# RNA at the surface of phase-separated condensates impacts their size and number

**DOI:** 10.1101/2021.06.22.449254

**Authors:** Audrey Cochard, Marina Garcia-Jove Navarro, Shunnichi Kashida, Michel Kress, Dominique Weil, Zoher Gueroui

## Abstract

Membrane-less organelles, by localizing and regulating complex biochemical reactions, are ubiquitous functional subunits of intracellular organization. They include a variety of nuclear and cytoplasmic ribonucleoprotein (RNP) condensates, such as nucleoli, P-bodies, germ granules and stress granules. While is it now recognized that specific RNA and protein families are critical for the biogenesis of RNP condensates, how these molecular constituents determine condensate size and morphology is unknown. To circumvent the biochemical complexity of endogenous RNP condensates, the use of programmable tools to reconstitute condensate formation with minimal constituents can be instrumental. Here we report a methodology to form RNA-containing condensates in living cells with controlled RNA and protein composition. Our bioengineered condensates are made of ArtiGranule scaffolds undergoing liquid-liquid phase separation in cells and programmed to specifically recruit a unique RNA species. We found that RNAs localized on condensate surface, either as isolated RNA molecules or as a homogenous corona of RNA molecules around the condensate. This simplified system allowed us to demonstrate that the size of the condensates scales with RNA surface density, the higher the RNA density is, the smaller and more frequent the condensates are. Our observations suggest a mechanism based on physical constraints, provided by RNAs localized on condensate surface, that limit condensate growth and coalescence.

---

It is increasingly recognized that biomolecular condensates contribute to organize cellular biochemistry by concentrating and compartmentalizing proteins and nucleic acids. They include a broad range of nuclear and cytoplasmic ribonucleoprotein (RNP) granules, such as nucleoli, P-bodies, germ granules and stress granules. Remarkably, abnormal condensate maturation into toxic aggregates is linked to viral infection, cancer, and neurodegenerative diseases^1^. Cellular condensates harbour a large diversity in terms of biochemical compositions as well as functions. Nevertheless, a unified model of formation via liquid-liquid phase separation (LLPS), where RNP constituents interact through multivalent and weak interactions, has been proposed to understand their biogenesis ^2–6^. In addition to their diverse compositions and functions, condensates are also diverse in size. Whereas P-bodies or PML bodies are often diffraction-limited puncta, other condensates such as germ granules, centrosomes, and nucleoli can reach few micrometres in size^2,7–10^. What sets condensate size and number in cells remains to be understood.

Mounting evidence based on *in vitro* reconstitutions and cellular approaches underlined the importance of multivalent interactions between RBPs and RNAs in shaping condensate biogenesis and morphology. In particular, RNA molecules have been proven to play fundamental roles in determining the structure, dynamic and biophysical properties of condensates^11^. For instance, RNAs act as molecular seeds to nucleate phase separated-condensates and regulate their assembly in a spatiotemporal manner^12–17^. On the opposite, high RNA concentration can dissolve condensates and keep prion-like RBPs soluble in cell nucleus^18,19^. In addition to their formation or dissolution, RNA molecules can also impact the viscosity of the RNP condensates as well as the dynamics of their components in a sequence-dependent manner^20–22^. The different structures of RNAs can determine the molecular specificity of RNP condensates and thus explain the coexistence of separate condensates with distinct molecular compositions^23^. Moreover, RNA-RNA interactions between unstructured RNAs can lead to the formation of non-spherical condensates^24^. Finally, RNAs can take part in RNA-RBP interactions that drive the formation of multiphasic condensates, whose structure relies on RNA concentration and on RNA-RBP interaction strength^22,25,26^. In addition to the contribution of condensate constituents, extrinsic factors such as membrane, cytoskeleton and chromatin can modulate LLPS and condensate biogenesis and coarsening^27–29^.

Interestingly, the biochemical and structural heterogeneity at the surface of condensates could also influence their stability. For instance in *C*.*elegans*, the adsorption of MEG-3 on PGL droplets drives the formation of a gel-like shell around a liquid core that eventually can stabilize P granules and trap RNAs^30,31^. Moreover, using an artificial scaffold in cells named ArtiGranule (ArtiG), we previously demonstrated that the condensation of the RNA-binding domain of the P-body protein Pumilio was sufficient to attract Pumilio RNA targets and P-body proteins at the surface of the condensates, which in turn impacted the seeding and the size of the condensates^32^. We therefore hypothesized that the size selection of the condensates relies on the adsorption of RNP elements at their surface, which may contribute to limit coarsening by steric exclusion^32^. However, extracting the specific role of RNA on condensate morphology is still inaccessible since natural RNP condensates emerge from a complex combination of RNA-protein, RNA-RNA, and protein-protein interactions^33–35^. To reduce this complexity, we developed a methodology to reconstitute the formation of RNA-containing condensates in living cells with controlled RNA and protein composition. Our bioengineered condensates were constituted of ArtiG scaffolds undergoing LLPS in cells, which were programmed to specifically recruit a unique RNA species. We first fused ArtiG scaffolds to an orthogonal RNA binding domain chosen to interact specifically with a synthetic mRNA. We found that RNAs localized on condensate surface, either as isolated RNA molecules or as a homogenous corona of RNA molecules around the condensate. The ArtiG condensates remained distinct from endogenous condensates, which enabled us to separate the role of the recruited RNA molecules from other potential contributors. We first observed a negative correlation between the number of condensates per cell and their mean diameter. By quantifying the localization and number of individual RNA molecules, we additionally found that the higher the RNA density is, the smaller and more numerous the condensates are. Overall, our data indicate that the size of RNP condensate scales with RNA surface density, which can be explained by physical constraints limiting condensate growth and coalescence.

## Reconstitution of RNA-protein condensates in human cells

Our first goal was to engineer artificial RNA-protein condensates that assemble through LLPS into cells. Our design combined two parts: a scaffold used to trigger the formation of protein condensates and a grafted RNA-binding domain to recruit specific RNA sequences (Fig. 1a). As protein scaffold, we used ArtiGs that form liquid protein condensates in a concentration-dependent manner through weak multivalent interactions^32^. The ArtiG scaffold developed previously was fused to a Pumilio-binding domain that recruits a large number of endogenous Pumilio RNA targets^32^. To restrict the targeting on ArtiGranules to one single RNA species, we chose an orthogonal RNA-binding domain, the MS2-coat protein (MCP), that recognizes specific MS2 stem-loops. The resulting plasmid construct, Fm-MCP-Ft, consisted in the fusion of an oligomeric ferritin (Ft) to MCP and a self-interacting domain F36M-FKBP (Fm) (Fig. 1a).

**Figure 1:**
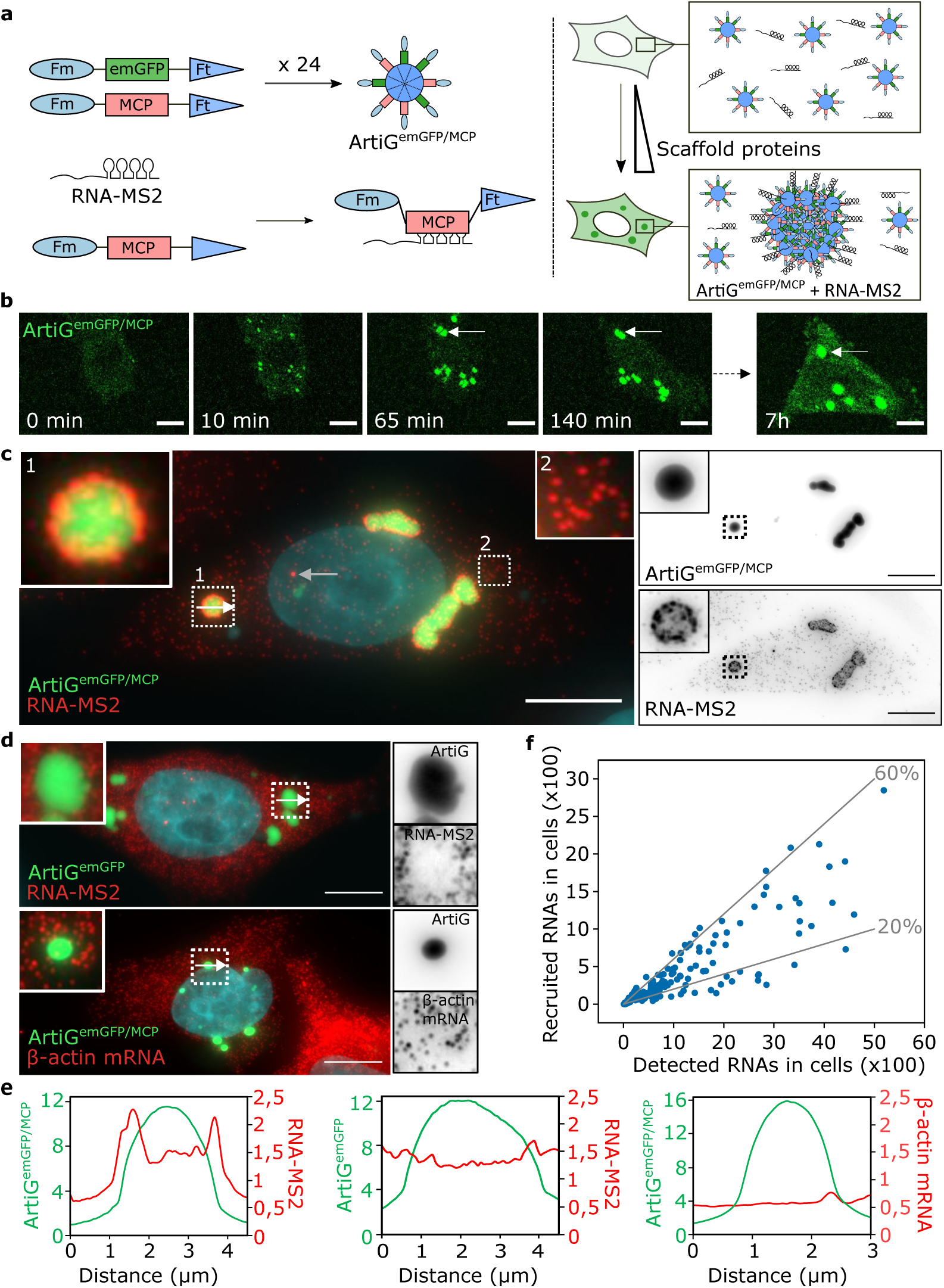
ArtiG^MCP^ condensates recruit a specific exogenous RNA. **a.** Schematic of the ArtiG^emGFP/MCP^ formation (Fm = F36M-FKBP, MCP = MS2 coat protein, Ft = Ferritin, ArtiG = ArtiGranule). ArtiG^emGFP/MCP^ form by LLPS driven by the homodimerization of Fm around Ferritin nanocages. MCP protein enable the recruitment of RNA-MS2 molecules to the condensates. **b.** Time-lapse confocal imaging of the formation of ArtiG^emGFP/MCP^ in Hela cells starting 8 hours after transfection with plasmids Fm-emGFP-Ft and Fm-MCP-Ft (1:1 plasmid ratio). The white arrow highlights a coalescence event. Scale = 10µm. **c.** Epifluorescence imaging of ArtiG^emGFP/MCP^ (green) and RNA-MS2 (red). Nuclei were stained with DAPI (blue in merge). The zoom in insert 1 shows the recruitment of RNA-MS2 around an ArtiG^emGFP/MCP^ condensate. Insert 2 shows isolated RNA-MS2 molecules. The white arrow indicates where the intensity profile in **e.** (left panel) was plotted. The grey arrow highlights a transcription focus. On the right panel, greyscale images correspond to separate channels. Scale = 10µm. **d.** Epifluorescence imaging of ArtiG^emGFP^ and RNA-MS2 (upper panel), and ArtiG^emGFP/MCP^ and β-actin mRNA (lower panel). The white arrows indicate where the intensity profiles in **e.** (middle and right panels) were plotted. Scale = 10µm. **e.** Intensity profiles across ArtiG condensates (white arrows in **c** and **d**). **f.** Number of RNA-MS2 molecules recruited at the surface of ArtiG^emGFP/MCP^ as a function of the total number of molecules detected in the cell, with each dot representing one cell (N = 140 from two independent experiments). Grey lines represent 20% and 60% of recruitment.

In order to monitor condensate formation in cells, we co-transfected HeLa cells with the multivalent MCP self-interacting scaffold Fm-MCP-Ft, and Fm-emGFP-Ft as a fluorescent tracer. Live confocal imaging performed 8 h after transfection showed that, initially, emGFP fluorescence at low expression level was diffuse in the cytoplasm. As Fm-emGFP-Ft expression increased, several bright fluorescent bodies nucleated throughout the cytoplasm and grew to reach a micrometric size within an hour (Fig. 1b). The emGFP expressing condensates, hereafter called ArtiG^emGFP/MCP^, were very mobile and rapidly grew as a function of time. When two proximal condensates docked, they tended to coalesce and to relax into large spherical bodies, as generally observed for endogenous liquid-like condensates (Fig. 1b, white arrow and Fig. S1a).

To reconstitute RNA-protein condensates using ArtiG^emGFP/MCP^, we first generated a plasmid to express a mRNA equipped with MS2 stem-loops in its 3’UTR (RNA-MS2, Fig. 1a). We co-transfected this plasmid and the plasmids expressing the ArtiG^emGFP/MCP^ scaffold and fixed the cells 24 h after transfection. We next monitored the intracellular localization of RNA-MS2 using single-molecule fluorescence in situ hybridization (smFISH)^36^. The majority of cells harboured micrometric ArtiG^emGFP/MCP^ condensates in the cytoplasm, with a striking Cy3-FISH signal around individual condensates, indicating a localization of RNA-MS2 molecules at the condensate surface (Fig. 1c, insert 1, and Fig. 1e, left panel). These RNAs were either present as isolated molecules or, when the number of recruited molecules was higher, were more homogeneously distributed around the condensates, into a corona made of a single RNA molecule layer (see other examples in Fig. S1b). Discrete Cy3 dots corresponding to individual mRNAs were also found dispersed throughout the cytosolic space (Fig. 1c, insert 2), as well as brighter spots in the nucleus corresponding to transcription foci (Fig. 1c, grey arrow). To assess the specificity of RNA recruitment on the ArtiG^emGFP/MCP^, we investigated the localization of RNA-MS2 in cells containing ArtiG^emGFP^ (devoid of the MCP domain) and found a complete absence of RNA-MS2 at the condensate periphery (Fig. 1d, upper panel, and Fig. 1e, middle panel). Experiments carried out in HEK293 cells showed the same results (Fig. S1c). Similarly, ArtiG^emGFP/MCP^ did not show any recruitment of the endogenous mRNA ß-actin and of the NORAD lncRNA (devoid of MS2) (Fig. 1d, lower panel, Fig. 1e, right panel, and Fig. S1d), thus confirming the specificity of the RNA recruitment. When quantifying the total number of mRNA molecules dispersed in the cytoplasm and localized on ArtiG^emGFP/MCP^, we found that 34% ± 19% of the cytoplasmic mRNAs were specifically recruited at the condensate surface (mean of 430 recruited RNAs and 1200 dispersed RNAs per cell) (Fig. 1f). Altogether, these data show that ArtiG^emGFP/MCP^ act as condensates localizing specific RNAs on their surfaces (ArtiG^emGFP/MCP/RNA^).

## Cytosolic RNAs are depleted at the vicinity of large condensates

As our data showed a robust recruitment of RNAs at the surface of ArtiG^emGFP/MCP^ condensates, we next investigated whether this recruitment impacted the distribution of RNAs in the cytoplasm. Interestingly, we observed a depletion of cytoplasmic MS2-RNAs close to ArtiG^emGFP/MCP/RNA^ condensates (Fig. 2a and Fig. S2a). This depletion was readily visible around large condensate clusters recruiting a high number of RNA molecules. On the examples shown in Fig. 2a and Fig. S2a, we quantified the density of RNAs as a function of the distance to the condensate edges. RNA density was almost zero in a large area ranging from the immediate vicinity of the cluster up to 3 µm from the condensate boarder. Then, RNA density increased until reaching a plateau at a distance of about 4 µm, with a value corresponding to the mean cytoplasmic RNA concentration found in cells (Fig. 2c, blue dots, and Fig. S2b). Likewise, plotting the cumulative number of RNAs outside the condensates as a function of the distance to the condensate edges showed first a very slow increase up to 3µm from the condensate edges (Fig. 2d, blue dots, and Fig S2c). Beyond this depletion area, the increase sharpened with a steady slope corresponding to an even cytoplasmic RNA concentration, except when the area occasionally included neighbouring condensates (Fig. 2c and d, light blue dots). For comparison, we quantified the spatial distribution of ß-actin mRNAs, which do not bind to ArtiG^MCP^. We found a total absence of depletion of ß-actin mRNAs around ArtiG clusters (Fig. 2b), with an even RNA density around the condensates (Fig. 2c and d, orange dots). Altogether, these results suggest that the RNA depletion was linked to the specific recruitment of RNA-MS2 on condensates and did not stem from potential non-specific steric exclusion around the condensates.

**Figure 2:**
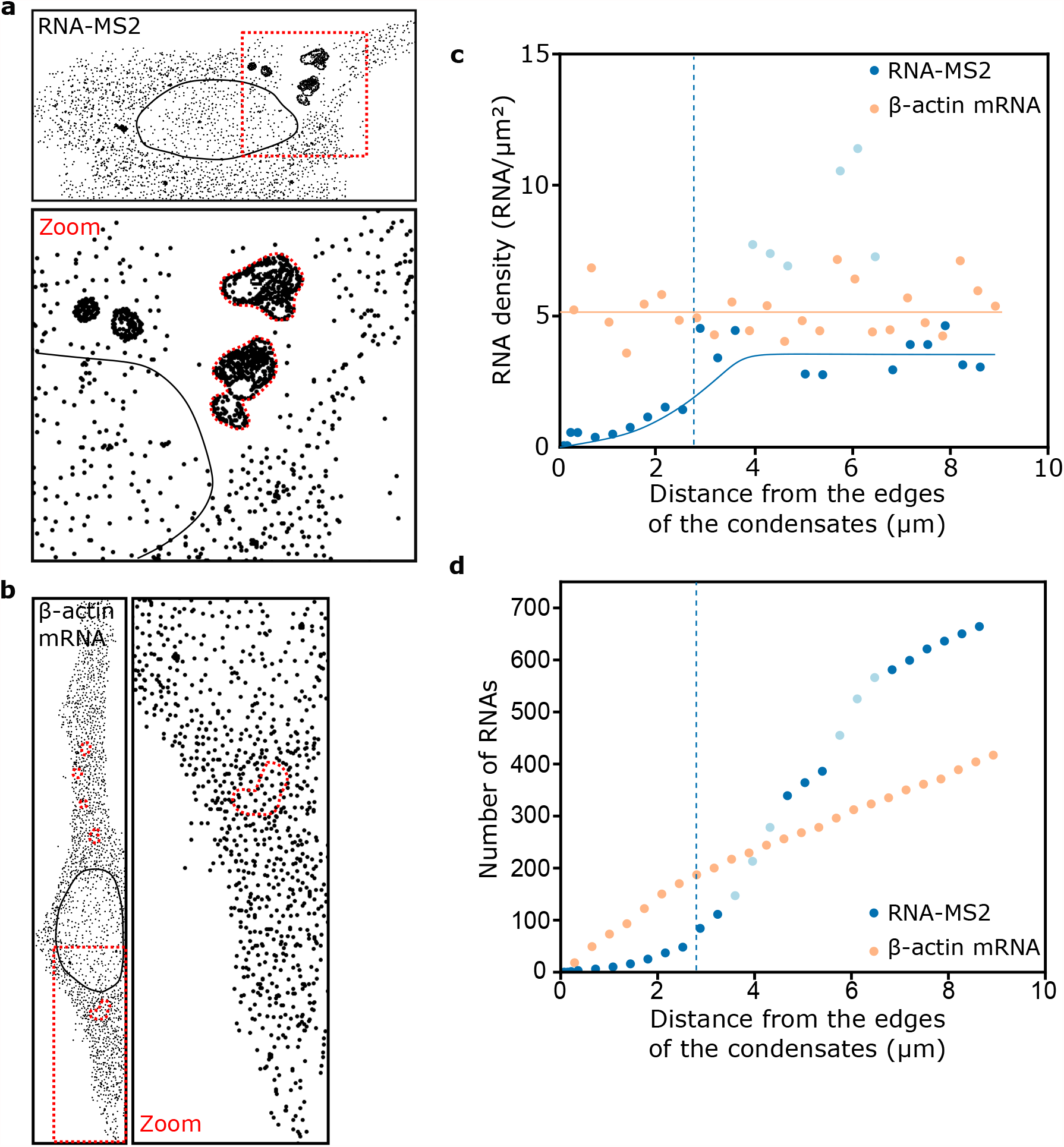
RNA-MS2 is depleted at the vicinity of ArtiG^emGFP/MCP^ condensates. **a.** RNA-MS2 molecules in a HeLa cell displaying a depletion of RNA-MS2 around ArtiG^emGFP/MCP^ condensates. The cooordinates of RNA-MS2 molecules have been acquired as described in the methods. Isolated dots are single RNA molecules while clustered dots overlap with ArtiG^emGFP/MCP^ condensates. The red square is enlarged below. **b.** β-actin mRNAs in a HeLa cell expressing ArtiG^emGFP/MCP^ (delineated in red). The red square is enlarged on the right. **c.** RNA-MS2 and β-actin mRNA density around the ArtiG^emGFP/MCP^ condensates delineated in red in **a.** and **b.** zoom panels. The orange line corresponds to β-actin mRNA mean concentration in the cytosol. The blue line corresponds to RNA-MS2 density that reaches a plateau after the depletion area. Dotted blue line marks the end of the depletion area. RNA densities when crossing neighbouring condensates are in light blue. **d.** Cumulative representation of the data shown in **c.**

## Artificial condensates are biochemically distinct from endogenous condensates

In a cellular context, biologically distinct RNP condensates that form in the same cytoplasm could interact with each other through shared proteins and RNAs, as described for P-bodies (PBs) and stress granules (SGs)^37^ or PBs and U-bodies^38^. Therefore, we next sought to investigate whether the local enrichment of mRNAs on ArtiG^emGFP/MCP^ may induce interactions with other cytoplasmic RNP granules. To this aim, we looked at the presence of PBs by immunostaining 24 h after transfection, using DDX6 as a PB marker. Our observations showed no particular physical proximity or docking between the two condensates (Fig. 3a, left panel). Similarly, there was not proximity between ArtiG^emGFP/MCP^ and SGs. Indeed, immunostaining of ATXN2L as a SG marker showed their absence 24 h after transfection (Fig. S3a), while no docking of ArtiG^emGFP/MCP/RNA^ with SGs was observed after SG induction with an arsenite stress (Fig. 3a, right panel). These results suggest that ArtiG^emGFP/MCP/RNA^ are biochemically distinct and physically independent from both PBs and SGs.

**Figure 3:**
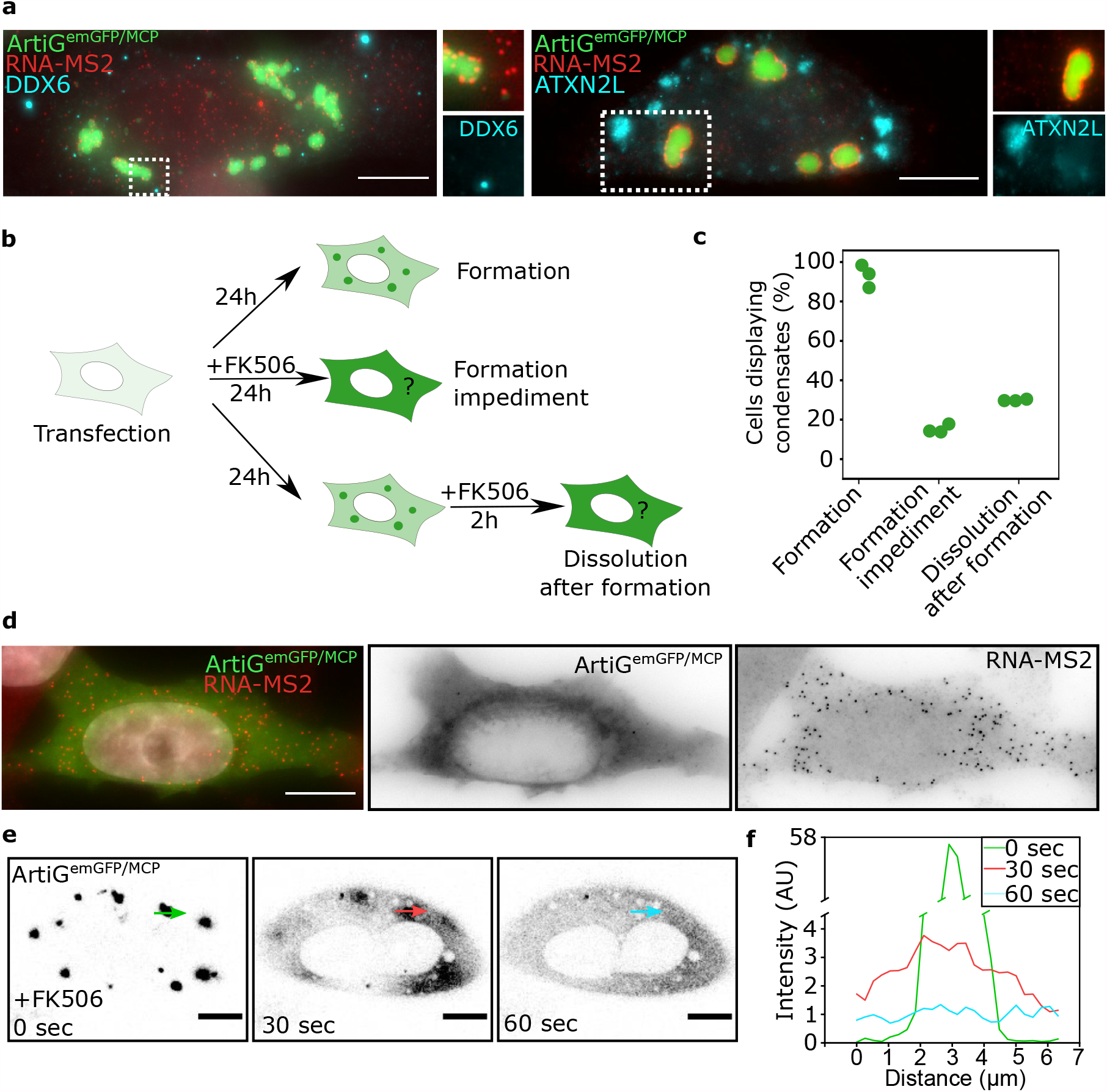
Interaction of ArtiG^emGFP/MCP^ with endogenous RNP granules, inhibition and reversibility. **a.** Epifluorescence imaging of ArtiG^emGFP/MCP^ (green) and RNA-MS2 (red) in HeLa cells, after immunostaining of DDX6 as a PB marker (left panel) or ATXN2L as a SG marker (right panel). In the right panel, SGs were induced with arsenite for 30 minutes. Scale = 10µm. **b.** Experimental design to test the inhibition and reversibility of ArtiG^emGFP/MCP^ formed in the presence of RNA-MS2, using FK506 in HeLa cells. FK506 was either added right after transfection to prevent condensation, or 24h after transfection to dissolve the condensates. **c.** Percentage of transfected cells displaying ArtiG^emGFP/MCP^ condensates in the absence of FK506 (left), when adding FK506 at the time of transfection (middle) or 24h later for 2h (right). **d.** Epifluorescence imaging of ArtiG^emGFP/MCP^ (green) and RNA-MS2 (red) after condensate dissolution with FK506. Nuclei were stained with DAPI (blue). Scale = 10µm. **e.** Confocal live imaging of ArtiG^emGFP/MCP^ dissolution. FK506 was added at time zero. Colored arrows indicate where the intensity profiles in **f.** were plotted. Scale = 10µm. **f.** EmGFP intensity profile across an ArtiG^emGFP/MCP^ condensate over time (colored arrows in **e.**).

## Controlled dissolution of artificial condensates

Recent studies suggest that the formation and stability of biological condensates are tightly regulated by multiple stimuli, including post-translational modifications, biochemical reactions, or physical parameters such as temperature or osmotic pressure changes^39,40^. By design, the formation and stability of the ArtiG^emGFP/MCP^ condensates are driven by multivalent interactions mediated by the Fm-Fm homodimer, and these interactions could in principle be disrupted by the addition of a chemical competitor, FK506. We therefore assessed if FK506 addition could first prevent condensate formation and secondly dissolve already formed condensates (Fig. 3b). In the absence of FK506, the majority of transfected cells exhibited ArtiG^emGFP/MCP/RNA^ condensates (93% after 24h of expression, Fig. 3c). This percentage dropped to 15% upon addition of FK506 at the time of transfection, with the majority of cells displaying a diffuse emGFP fluorescence and a homogeneous MS2-RNA distribution in the cytosol (Fig. 3d). Thus, FK506 efficiently inhibited the formation of the condensates. In a second experimental design, we examined the dissolution of fully formed ArtiGs by adding FK506 24h after transfection (Fig. 3b). After 2h of FK506 incubation, we found that the majority of cells lacked ArtiGs and displayed diffuse emGFP with Cy3-labelled MS2-RNAs distributed throughout the cytoplasm (70%, Fig. 3c). Thus, FK506 treatment induced the dissolution of the majority of pre-formed ArtiG^emGFP/MCP/RNA^ condensates. To further characterize FK506 effect, we examined the time-scale of dissolution using live confocal microscopy. Upon addition of FK506, some cells exhibited condensates dissolving within few seconds (Fig. 3e and supplementary video 2), while in others dissolution took up to 30 minutes (Fig. S3b). These dissolutions were accompanied with a strong increase of the cytosolic fluorescence signal, corresponding to the release of the ArtiG scaffold (Fig. 3e). We also observed a few cells with smaller condensates and a stronger cytosolic fluorescence, corresponding to incomplete condensate dissolution, in agreement with the observation of residual condensates in fixed cells (Fig. S3c). Altogether, our data showed that pre-treatment with the FK506 binding competitor of Fm proteins provides a mean of preventing the formation of the ArtiG condensates, while it globally induces their disassembly when they are already formed. Our system thus allows for a controlled inhibition and disassembly (by adding FK506) of artificial condensates in living cells.

## Linking condensate size and number of recruited RNAs

Determining how the primary constituents of condensates set the variety of condensate size and morphology naturally observed in cells remains very complex, since RNAs and proteins establish a large network of interactions. The ArtiG condensates potentially provide an important simplification to this problem, as only one RNA species is recruited to the condensates. We could therefore analyse how RNA contributes to ArtiG^emGFP/MCP/RNA^ condensate morphology, by quantifying the recruitment of RNAs in condensates and condensate size within single cells. The size and number of ArtiG^emGFP/MCP/RNA^ condensates were heterogenous between cells, with cells exhibiting few condensates and others a larger number (Fig. 4a). The distribution of the mean diameter of ArtiG condensates roughly ranged from 0.4 to 4 µm depending on the cell (mean ± SD = 1.1 ± 0.6 µm, coefficient of variation CV = 58%, Fig. 4b, left panel). While 75% of the cells had condensates with a mean diameter below 1.5 µm, we observed particularly large condensates, up to 4 µm in diameter, in the other cells. The number of condensates per cell was also very diverse (mean number = 33, CV = 116%, Fig. 4b, right panel). Interestingly, condensate mean size per cell was inversely proportional to their number (Fig. 4c). Indeed, cells displaying large condensates (> 1.5µm of diameter) always had a limited number of them (<25). In contrast, a higher number of condensates in a cell (>25) was correlated with a mean diameter of the condensates below 1.5 µm. As opposed to the heterogeneity of condensate size between cells, we found a homogeneity of size within a given cell (average CV=30%, Fig. 4d).

**Figure 4:**
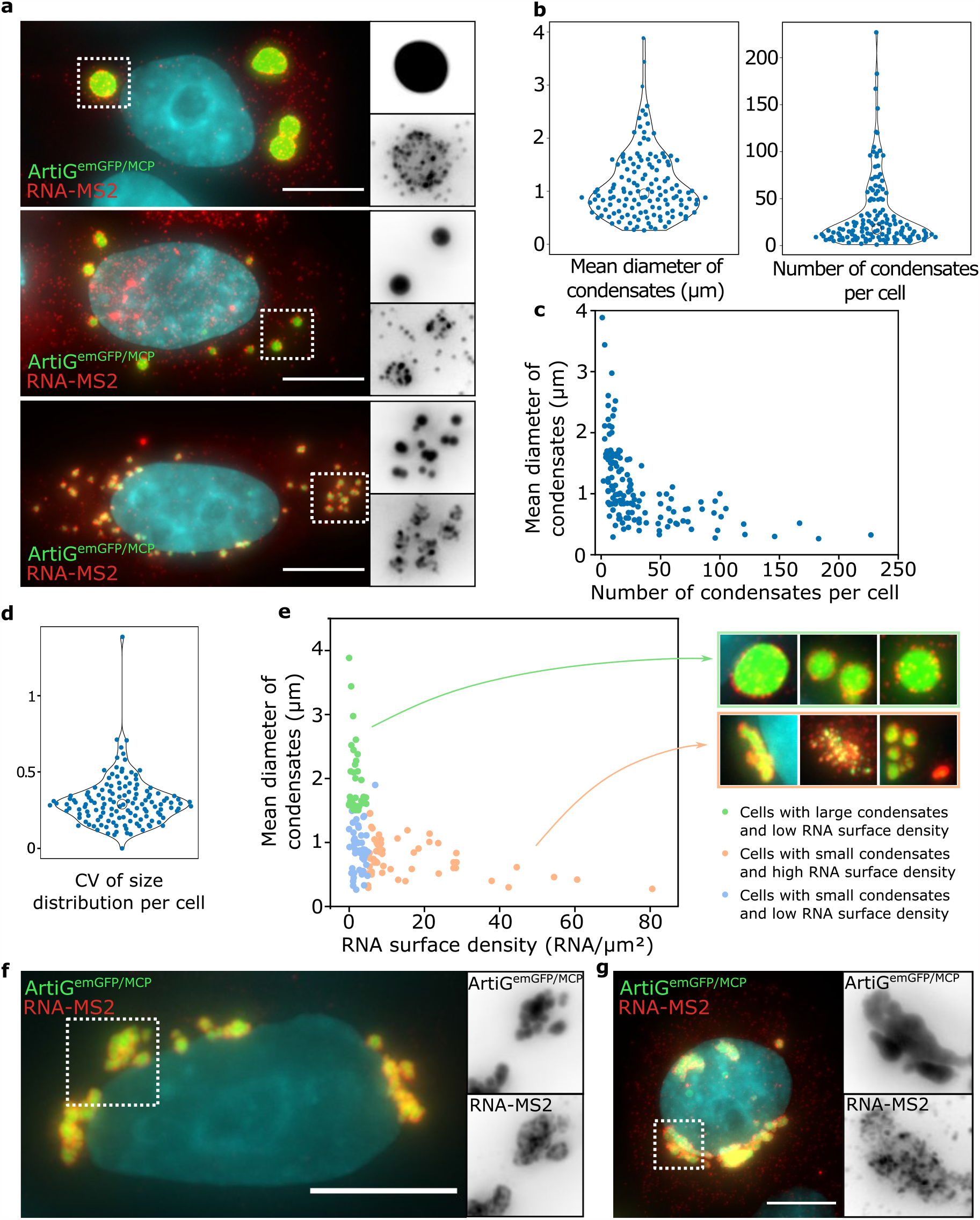
Heterogeneity in the morphology of ArtiG^emGFP/MCP^ condensates. **a.** Epifluorescence imaging of three Hela cells displaying different sizes and numbers of ArtiG^emGFP/MCP^ in the presence of RNA-MS2. Scale = 10µm. **b.** Distribution of the mean diameter of condensates per cell (left panel) and the number of condensates per cell (right panel), with each dot representing one cell (N = 140 from two independent experiments). **c.** Mean diameter of the condensates as a function of the number of condensates. **d.** Distribution of the coefficients of variation (CV) of the size distribution. **e.** Mean diameter of the condensates as a function of RNA surface density. Green and orange dots highlight cells displaying a mean diameter above or below 1.5µm, respectively, and an RNA surface density below or above 5 molecules/ µm ^2^respectively. Images on the right correspond to examples of condensates for the green and orange cell categories. **f. g.** Example of well-defined **(f.)** and intertwining **(g.)** condensate clusters.

We next sought to examine whether there was a correlation between condensate number and size, and RNA recruitment. To this aim, we computed, per cell, the density of RNAs recruited at the surface of ArtiG condensates and their mean diameter (Fig. 4e). We could highlight two groups of cells. In cells displaying larger condensates (mean diameter > 1.5 µm, mean ± SD = 2.0 ± 0.6 µm), the RNA surface density was below 5 RNAs/µm^2^ (mean = 2.0 RNAs/µm^2^, Fig. 4e, green dots). These condensates were generally spherical with a small number of RNAs at their periphery. In contrast, a higher RNA density (> 5 RNAs/µm^2^, mean = 16.0 RNA/µm^2^) was correlated with a mean diameter of ArtiG below 1.5 µm (mean ± SD = 0.79 ± 0.32 µm, Fig. 4e, orange dots). In these cells, condensates were often found in close proximity to each other, forming cluster-like patterns (more than 5 condensates docking together) with RNA patches or corona separating individual condensates (Fig. 4f). These clusters were reminiscent of coalescence events but their high number suggested that the coalescence process was arrested, so that condensates did not relax into a sphere. We even observed a few cases where ArtiG^emGFP/MCP^ and RNA molecules seemed intertwined, with frontiers between condensates becoming blurred and ArtiG^emGFP/MCP/RNA^ losing their round shape (Fig. 4g). To sum up, we found that all large spherical ArtiG condensates displayed few RNA molecules on their surfaces, while condensates with a high RNA surface density had a smaller size.

## Evolution of condensate size and morphology as a function of RNA surface density

To refine our analysis of the role of RNA localization in the condensate morphology, we increased the expression of transcribed RNA-MS2 by transfecting a larger quantity of plasmids in cells (five-fold more). In these conditions (RNA high) the mean number of RNA-MS2 transcripts per cell rose from about 1200 to around 2400, and the mean number of RNA-MS2 recruited at the surface of the condensates from 430 to 1100. This RNA high condition did not modify the expression of the ArtiG scaffold, as indicated by Western blotting experiments (Fig. S5). Firstly, we found that experiments performed in RNA high condition led to smaller condensate size (0.70 ± 0.32 µm instead of 1.25 ± 0.69 µm) (Fig. 5a), which supports our observation that higher RNA surface localization resulted in smaller condensates. Furthermore, in the RNA high condition, almost no cells displayed large condensates (diameter > 1.5µm) (Fig. 5b) and a higher incidence of the cluster-like patterns, about 58 % of cells, was observed in regard to 42.5 % in the RNA low condition (Fig. 5c). Among the clusters, condensate intertwining, which was a rare event in the low RNA condition (5 %), became more common (17 %). Altogether these data confirm a direct relationship between RNA surface density and condensate size and number, and underlie that the average size of ArtiG condensates is influenced by the level of expression of the recruited RNA.

**Figure 5:**
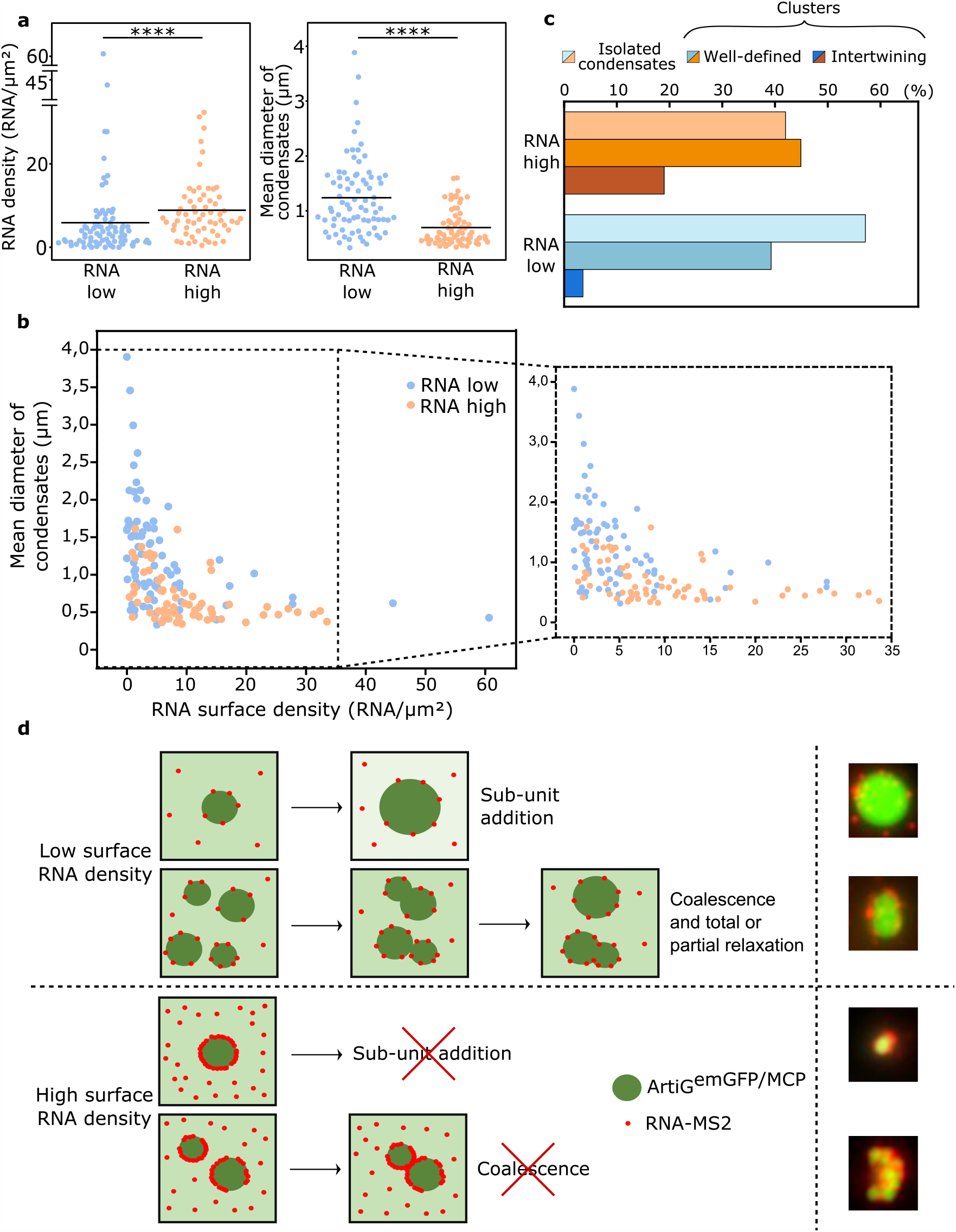
Impact of RNA recruitment at the surface of condensates. **a.** Distribution of RNA-MS2 density at the surface of condensates (left panel) and of mean diameter of the condensates (right panel) in RNA low and RNA high conditions, with each dot representing one cell (N = 80 for RNA low and N = 69 for RNA high conditions, both from two independent experiments). Differences between RNA low and RNA high conditions were statistically significant using a Wilcoxon rank sum test: p-values < 10-5. **b.** Mean diameter of ArtiG^emGFP/MCP^ in cells as a function of the RNA-MS2 surface density in RNA low (blue dots) and RNA high (orange dots) conditions. The area of the graph framed in black was enlarged in the right panel. **c.** Percentage of cells with ArtiG^emGFP/McP^ that display isolated condensates, clusters of well-defined condensates, or intertwining condensates, in low RNA and high RNA conditions. **d.** Schematic model for the impact of surface RNA molecules on condensate growth and coalescence. Illustrative examples of ArtiG^emGFP/McP^ (in green) and RNA-MS2 (in red) epifluorescence images are shown on the right.

## Discussion

RNA is more and more recognized as a driving force in cellular organization and functions. These polymers can interact and scaffold hundreds of proteins to generate high order organizations including RNP condensates. The first high throughput biochemical studies of RNP condensates showed that their RNA and protein content is highly complex^41–44^. While these studies highlight that condensates assembly is driven by the combination of multiple RNA-protein, protein-protein, and RNA-RNA interactions, the rules governing RNA and protein spatiotemporal co-assembly are still enigmatic. As a consequence, deciphering and manipulating RNP condensates *in cellulo* still remains a difficult task. In this context, *in vitro* reconstitutions using purified components have been a powerful strategy to study the diverse roles of RNAs involved in specifying the structure, composition and dynamical properties of RNP granules^11,22,25,45,46^. Physicochemical parameters defining RNA polymers such as their length, chemical complexity and sequence, could thus be assessed in a reconstituted environment. Despite their obvious advantages, several limitations arose from these reductionist approaches. For instance, the physiological relevance of the concentration and composition of *in vitro* model condensates can be questioned, as well as their environment that lacks the physicochemical complexity present within cells. Alternatively, the over-expression of proteins, often chosen among scaffolding proteins founded to drive LLPS in test tube, was also widely used to identify the propensity of specific protein domains to undergo phase separation in a cellular environment^19,47,48^.

To combine the level of controls provided by *in vitro* experiments with the possibility to study LLPS within the cellular environment, several bioengineering approaches have been developed to form, within cells, condensates with specific properties. These approaches often use optogenetic and chemical actuations based on protein-multimerization domains acting as scaffolds of artificial condensates. Being able to engineer condensates in cells with well-defined compositions, structures and dynamical properties provides novel insights to correlate condensates biochemical functions with their material states, a link that is still difficult to reach using conventional cell biology techniques. Recent developments and applications of such engineered condensates have thus enabled the quantitative studies of the dynamical properties of phase-separated condensates within cytoplasm and nucleus. For example, light-induced strategies based on optoDroplets allowed the actuation of model condensates mimicking pathological assemblies appearing in some age-related diseases^49,50^. Alternative synthetic protein condensates were designed with programmable material properties or functions^51–58^. In order to reconstitute artificial RNP condensates, iPOLYMER and ArtiGs were designed to induce condensation of RNA-binding motifs found in SGs (TIA-1) and PBs (Pumilio), respectively^32,59^. While these approaches provided a powerful mean to manipulate RNP condensate mimics in cells, the use of motifs from RBPs that are known to target thousands of RNA species could limit the understanding of observed effect in cells. To overcome this limitation, our approach was to reconstitute in cells the formation of artificial RNP condensates using ArtiGs modified to target a single RNA species. Remarkably, recruited RNAs localized at the condensate surface and bound as isolated RNA molecules or as homogenous RNA coronas surrounding the condensates (Fig. 1 and S1).

How these heterogenous patchy or corona-like patterns emerge from the interactions between ArtiG scaffolds and MS2-RNAs? Several *in vitro* studies and numerical simulations reported how multilayered organizations, such as core-shell droplets, assembled from ternary systems composed of protein-RNA interacting molecules^22,25,26,60–62^. A possible mode of formation of these multiphase droplets results from competing intermolecular interactions between macromolecular constituents that drive differences in surface tension and coexisting liquid phases. In this regard, our RNP condensates differ from co-existing liquid phases that demix into core-shell droplets, since they generally displayed a single RNA molecule layer as a shell covering ArtiG condensates. Instead, the assembly pathway controlling the formation of condensates with an RNA corona could arise from a stepwise process: first, ArtiG^MCP^ scaffolds undergo LLPS and, subsequently, RNA molecules are recruited on the condensate surface with an efficiency that depends on the RNA expression level (Fig 5d).

The robust formation of ArtiG condensates in cells provides an efficient assay to examine basic questions such as how condensate size scales with RNA surface density. The ability to quantify individual RNA molecules on ArtiG condensate surface provides a unique mean to establish a direct link between RNA surface localization and condensate size and number. We found that the RNA density at the surface of condensates was correlated to their size and number, with large condensates displaying only a few RNAs on their surface whereas high RNA density always implied smaller and more numerous condensates. Furthermore, when we increased RNA expression in cells, and consequently RNA surface density on condensates, cells harboured smaller condensates, which supports a causal relationship between RNA surface density and condensate size.

Several examples in cell biology suggest the existence of a scaling of cellular organelles with cell volume, which could be understood if cells contain a limiting pool of structural components supporting the organelle assembly^9,63,64^. In cells, native condensates such as PBs and PML nuclear bodies are generally found as sub-micrometric bodies that often do not grow over a certain size. This is generally in contradiction with the thermodynamical equilibrium picture of phase separated systems predicting an evolution towards a single condensed phase co-existing with a dilute phase. Initial growth of phase-separated condensates generally occurs through subunit addition, and coarsening through coalescence or Ostwald ripening^65–67^. Thus, a solution to regulate condensate size would be to tune one of those three pathways (subunit addition, coalescence and Ostwald ripening), either through physicochemical parameters or by modifying interaction strengths and valences by biochemical reactions such as post-translational modifications^68,69^. Recent theoretical studies suggest that both active and passive processes can be in play^70,71^. For instance, it has been proposed that active processes within condensates could suppress Ostwald ripening and account for size selection^63,72–75^. However, in the case of ArtiG^emGFP/MCP/RNA^, the two main formation pathways are subunit addition and coalescence (Fig. 1b). Client proteins acting like surfactants may reduce the energy required for the formation of an interface between the dense and dilute phase and lead to size-conserved multi-droplet systems instead of the expected single large condensed phase, with condensate size decreasing as a result of an increase in the client concentration.^76^ In this respect, the protein Ki-67, localized at the surface of chromosomes, may for instance form a steric barrier that prevents the chromosomes from collapsing into a single entity^77^. A high surface charge density and thus a high electrostatic repulsion between biomolecular condensates may alter their propensity to fuse^78^. *In vitro* observations of the co-assembly between RNA homopolymers and mRNAs showed multi-phase assemblies, with RNAs localized at the droplet surface, suggesting that RNAs act as an interfacial shell stabilizing multiphasic condensates^45^. Alternatively, a recent study explained how the RNA shell-forming domain of paraspeckles can modulate condensate shape and size and suggested a micellization-based model of assembly^79^. Here we propose that the RNA present at the surface of ArtiG condensates cause a steric hindrance that may prevent the growth of condensates by both subunit addition and coalescence (Fig. 5d). In this picture, RNAs could contribute to the colloidal stability of the condensates and thus regulate their size and number.

At a high RNA surface density, we found that ArtiG condensates can lose their sphericity and adopt a clustered morphology reminiscent of TIS granules^24^. In contrast to TIS granules, where a skeleton made of RNA-RNA interactions between unstructured regions counterbalances the excess of surface energy generally driving fusion and relaxation, here RNAs at the condensate surface could impede coalescence by steric hindrance. We could also envision the existence of intermolecular interactions between RNAs that would bridge adjacent ArtiG condensates and consolidate their stability.

Such an impact of surface RNA on condensate morphology may be relevant for native RNP condensates. Indeed, the spatiotemporal organization of RNAs at the surface of native condensates has recently been investigated using advanced imaging tools. For example, super-resolution imaging showed that RNAs and proteins composing SGs are enriched within a main solid core surrounded by a less concentred layer^44,80^. It has also been shown that RNAs exhibit diverse localization within PB core and at their surface^81^. On SG or PB surface, RNAs can make transient contacts before stably associating inside the granules or leaving the granules for an alternative fate^81,82^. Some of these RNAs are coding mRNAs, thus associating with ribosomes and other translation-related proteins^83^, while others are long non-coding RNAs with a regulatory function^81^. These RNAs can partition bidirectionally between biologically different condensates^82,84^. In the case of germ PBs, the association of the RNAs with the surface of the condensates can even be required for translation to happen^85^. Along this line, our work suggests that localization of RNAs at the condensate surface could also feedback on condensate biogenesis.

In conclusion, our methodology to reconstitute biomolecular condensates in cells with controlled compositions and properties has proved powerful to reveal the role of RNA in condensate morphology. Not only its flexibility will enable to address other such basic biological issues in the future, but it could also be a mean to engineer novel properties within cells. More generally, our study stresses the importance of an understudied aspect of condensates, which is the role of the biomolecules present at their surface. It illustrates how chemical and physical heterogeneities on condensate surface may determine RNA granule morphogenetic properties.

## MATERIALS AND METHODS

### Experimental model

Human epithelioid carcinoma HeLa (ATCC, ccl-2) and embryonic kidney HEK-293 (ATCC, CRL-1573) cells were maintained in Dulbecco’s modified Eagle’s medium (with 4.5 g/L D-glucose, HyClone) supplemented with 10% fetal bovine serum (PAA Laboratories) and antibiotics, at 37 °C in a 5% CO2 humidified atmosphere. Cells were routinely tested for mycoplasma contamination. Stress granules were induced with 0.5 mM sodium arsenite (Sigma) for 30 min at 37°C. For inhibition and reversibility experiments (Fig. 3b-e), 2.5 µM of FK506 was added to the cell culture medium.

### Plasmids

All constructs were sub-cloned into pcDNA 3.1 plasmid (Invitrogen). The plasmids pcDNA3.1-F36M-FKBP(Fm)-emGFP-hFt and pcDNA3.1–Fm–hFt were previously described^32^. The plasmid pcDNA3.1–Fm–MCP–hFt was obtained by replacing the emGFP CDS between XhoI and BamHI restriction sites in pcDNA3.1–Fm–emGFP–hFt by a CDS encoding a tandem of two MS2-coat proteins (MCP). We used a tandem to improve the efficiency of binding to MS2 stem loops^86^. RNA-MS2 was expressed from the plasmid pcDNA3.1–4xMS2, which was obtained by inserting the iRFP CDS in the pcDNA3.1 backbone and inserting a tandem of four MS2 stem loops in the 3’UTR.

### Transfection

For live experiments, HeLa cells were cultured on 35mm µ-dishes with polymer coverslip bottom (Ibidi). For other experiments, HeLa cells and HEK-293 cells were cultured on 22×22mm glass coverslips (VWR) in 6-well plates (Falcon). 24h after seeding, transient transfection using Lipofectamine 2000 (Invitrogen) was carried out according to the manufacturer’s protocol. For live experiments cells were transfected with a 1:1 ratio of pcDNA3.1–Fm–MCP–hFt and pcDNA3.1– Fm–emGF –hFt (800ng total per Ibidi) and 20ng of pcDNA3.1– 4xMS2. For other experiments, cells were transfected with a 1:0.7:0.3 ratio of pcDNA3.1–Fm–MCP–hFt, pcDNA3.1–Fm–emGFP–hFt and pcDNA3.1–Fm–hFt (2µg total per well) and 50ng (low RNA), 250ng (high RNA) or indicated amount (Fig. S4) of pcDNA3.1– 4xMS2.

### Single molecule Fluorescence In Situ Hybridization (smFISH)

Single RNA molecule detection was performed according to the previously described smiFISH method^36^. 24h after transfection, cells were fixed in 4% paraformaldehyde (PFA) for 20 min at RT, and permeabilized with 70% Ethanol in phosphate buffer saline (PBS) at 4°C overnight. For each target RNA (RNA-MS2, β-actin mRNA and NORAD lncRNA), a set of 24 primary probes, constituted of a sequence-specific part and a common FLAP sequence (TTACACTCGGACCTCGTCGACATGCATT), were designed with the Oligostan R script^36^. Gene-specific probes and the Cy3 FLAP probe (sequences in the Supplementary table 1) were purchased from Integrated DNA Technologies.

### Immunofluorescence

At the end of the smFISH steps, cells were permeabilized with Triton X-100 0.1% in PBS for 10 min at RT, washed twice with PBS at RT, incubated with the primary antibody, washed three times with PBS at RT for 5 min, incubated with the secondary antibody, washed three times with PBS at RT for 5 min and finally mounted with VECTASHIELD mounting medium (Vector Laboratories). Primary antibodies were rabbit antibodies against DDX6 (Bethyl, 1:1000 dilution) and rabbit antibodies against ATXN2L (Bethyl, 1:500 dilution), diluted in PBS 0.1% BSA. The secondary antibody was F(ab’)2-Goat anti-Rabbit IgG conjugated with Alexa Fluor 350 dye (Invitrogen, 1:500 dilution).

### Imaging

For live experiments, cells were imaged on a Zeiss LSM 710 META laser scanning confocal microscope using an ×63 oil-immersion objective (PlanApochromatic, numerical aperture (NA) 1.4), at 37 °C in a 5% CO2 humidified atmosphere, either starting 8h after transfection (ArtiG formation) or 24h after transfection (ArtiG dissolution). Microscope hardware and image acquisition were controlled with LSM Software Zen 2009. Images were analyzed using ImageJ (NIH).

For smFISH experiments, cells were imaged by epifluorescence microscopy performed on an inverted Zeiss Z1 microscope equipped with a motorized stage using a ×63 (NA 1.32) oil-immersion objective. Images were processed with open-source software Fiji and Icy^87,88^.

### Western Blotting

24h after transfection, cells were collected using trypsin (Gibco) and resuspended in Laemmli 1X buffer. Proteins were denatured at 100°C for 5 min. Soluble proteins were retrieved after centrifugation at 15000 g at 4°C for 10 minutes and quantified using the Coomassie protein assay (Thermo Scientific). 25 µg of proteins were separated on a NuPAGE 4% -12% gel (Invitrogen, Thermo Fisher Scientific) and transferred to an Optitran BA-S83 nitrocellulose membrane (GE Healthcare Life Science). The membrane was then blocked in PBS with 5% non-fat milk for 30 min, before being incubated with the primary antibody (6x-His Tag Monoclonal Antibody, Invitrogen) overnight at 4°C. After washing the membrane five times for 5 min with PBS, it was incubated for 30 min in PBS with 5% non-fat milk and then for 1h at RT with horseradish peroxidase-conjugated secondary anti-mouse antibody (1:10000 dilution in PBS with 5% non-fat milk, Jackson Immunoresearch Laboratories), and washed again. Proteins were detected with the chemiluminescence detection reagent Perkin Western Lightning plus ECL (Perkin Elmer) and visualized using a radiology film processor (Curix 60, AGFA).

### Data analysis

Detection and counting of RNA molecules was performed using version 0.4.0 of Python package Big-FISH (https://github.com/fish-quant/big-fish). A binary mask was created on the emGFP channel to detect the ArtiG^emGFP/MCP^ condensates. To count the number of recruited RNAs in individual cells, all RNA molecules were detected, and their coordinates were compared to the binary mask. For the analysis of RNA depletion at the vicinity of condensates, the binary mask was repeatedly expanded by 5 pixels and RNA molecules in the mask were counted at each step, which enabled the calculation of both the number and the density of RNA molecules in the last incremented area. For the quantification of the correlation between condensates size and surface RNA density, condensates sizes were measured using Icy spot detector^89^. When close condensates were not discriminated, the detected regions of interest were adjusted by hand. Clusters that were impossible to separate into individual condensates were excluded from the statistics of size. For consistency, the few large clusters included in the statistics of Figure 4 (low RNA condition) were excluded in the statistics of Figure 5, and large clusters in the high RNA condition were similarly excluded. The exterior surface was calculated for each condensate from the surface of the condensate’s maximum projection. Then, for each cell, the sum of the surface of all condensates was calculated and used to determine the mean RNA density at the condensate surface (ratio of the total recruited RNAs to the sum of the condensate surface).

Formatting of cell images was performed using the open-source software Fiji^87^. For Figure 2 and S2, the RNA coordinates were first saved from the Python workflow and then opened on Fiji. Graphic were generated using the shiny app PlotsofData^90^ (violin plots in Figure 4b, 4d and 5a) and OriginPro (OriginLab). For all violin plots, circles correspond to the mean. Schema (Fig. 1a and 5d) were drawn with the open-source vector graphics editor Inkscape. For Figure 5a, Wilcoxon rank-sum tests (nonparametric test to compare two distributions) were performed using the *ranksum* Matlab function (MathWorks).

## Acknowledgments

The authors acknowledge all the members of the Biophysical Chemistry group of the École normale supérieure for fruitful discussion. We thank A. Imbert, M.A Plamont, M. Ernoult-Lange, M.N Benassy, M. Bénard, H. Saito, and S. Matsumoto for their help along the project, as well as Adham Safieddine for carefully reading the manuscript. A.C was supported by IPV-SU PhD fellowship. M.G.J.N was supported by FRM (ING20150532742). This work was supported by the CNRS, Ecole Normale Supérieure, and ARC (20181208003) to Z.G., FRM (MND202003011470) and iBio (SU) to Z.G and D.W, and ANR (ANR-19-CE12-0024-01) to D.W (ANR-19-CE12-0024-01).

## Author Contributions

A.C, D.W and Z.G conceived the project and analyzed results. A.C carried out and analyzed most experiments. M.G.J.N contributed to the design of protein scaffolds and performed initial experiments. S.K and M.K contributed to the design of synthetic RNAs. A.C, D.W and Z.G wrote the manuscript and all authors were involved in revising it critically for important intellectual content.

## Declaration of Interests

The authors declare that they have no competing interests.

## Supporting Information

4 supplementary figures and 1 supplementary table.

---

**Figure S1:**
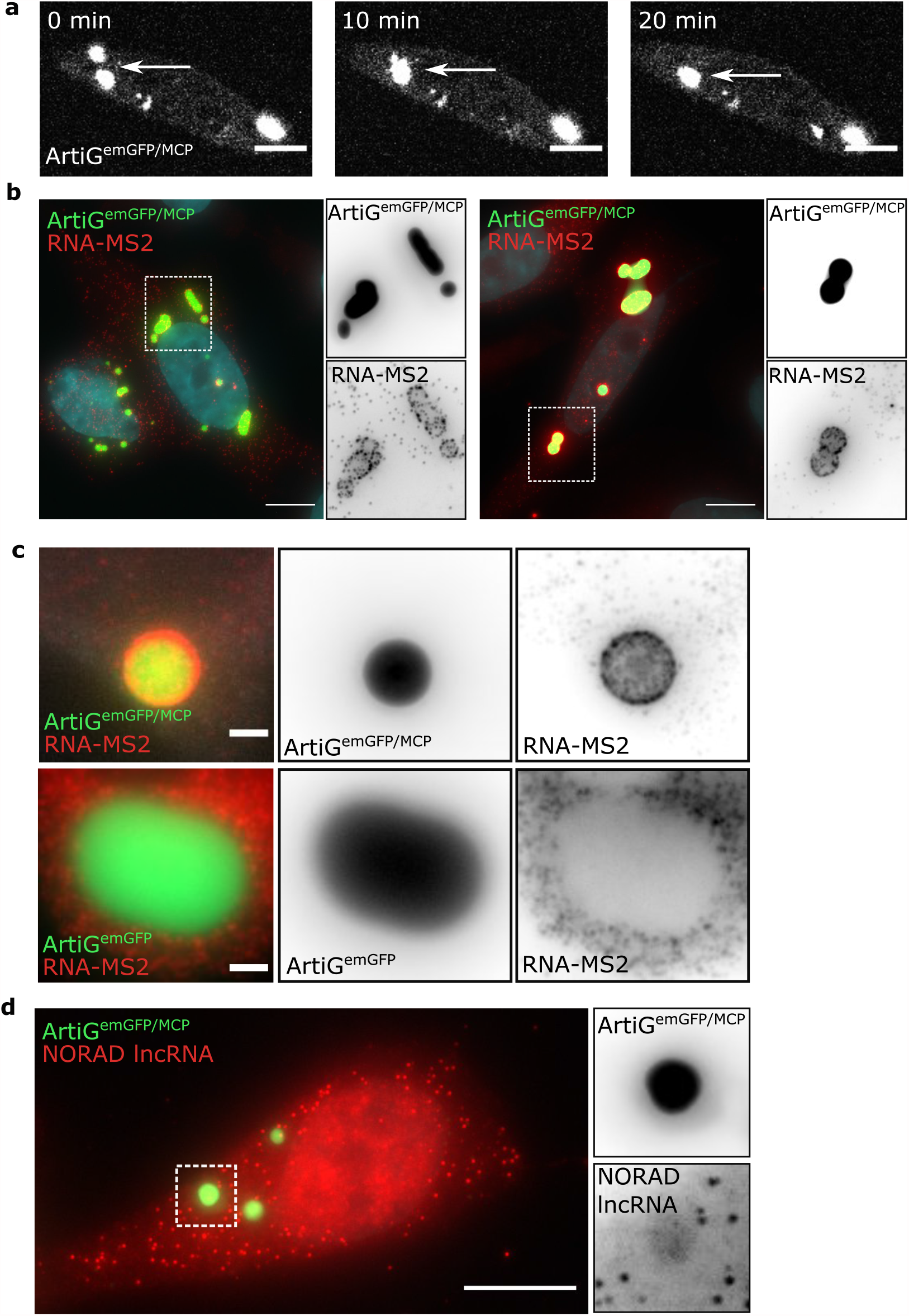
Validation of ArtiG^emGFP/MCP^ specificity. **a.** Coalescence of two ArtiG^emGFP/MCP^ condensates in a Hela cell and relaxation into a spherical shape (white arrow). Scale bar = l0µm. **b.** Epifluorescence imaging of two Hela cells displaying the usual surface recruitment of MS2-RNA (red) around ArtiG^emGFP/McP^ (green). Greyscale images show zooms of separate channels. Scale bars = l0µm. **c.** Epifluorescence imaging of one ArtiG^emGFP/McP^ (upper panel) and one ArtiG^emGFP^ (lower panel) and RNA-MS2 in HEK293 cells. Scale bars = 2µm. **d.** Epifluorescence imaging of ArtiG^emGFP/MCP^ (green) and NORAD lncRNA (red) in Hela cells. Scale bar = l0µm.

**Figure S2:**
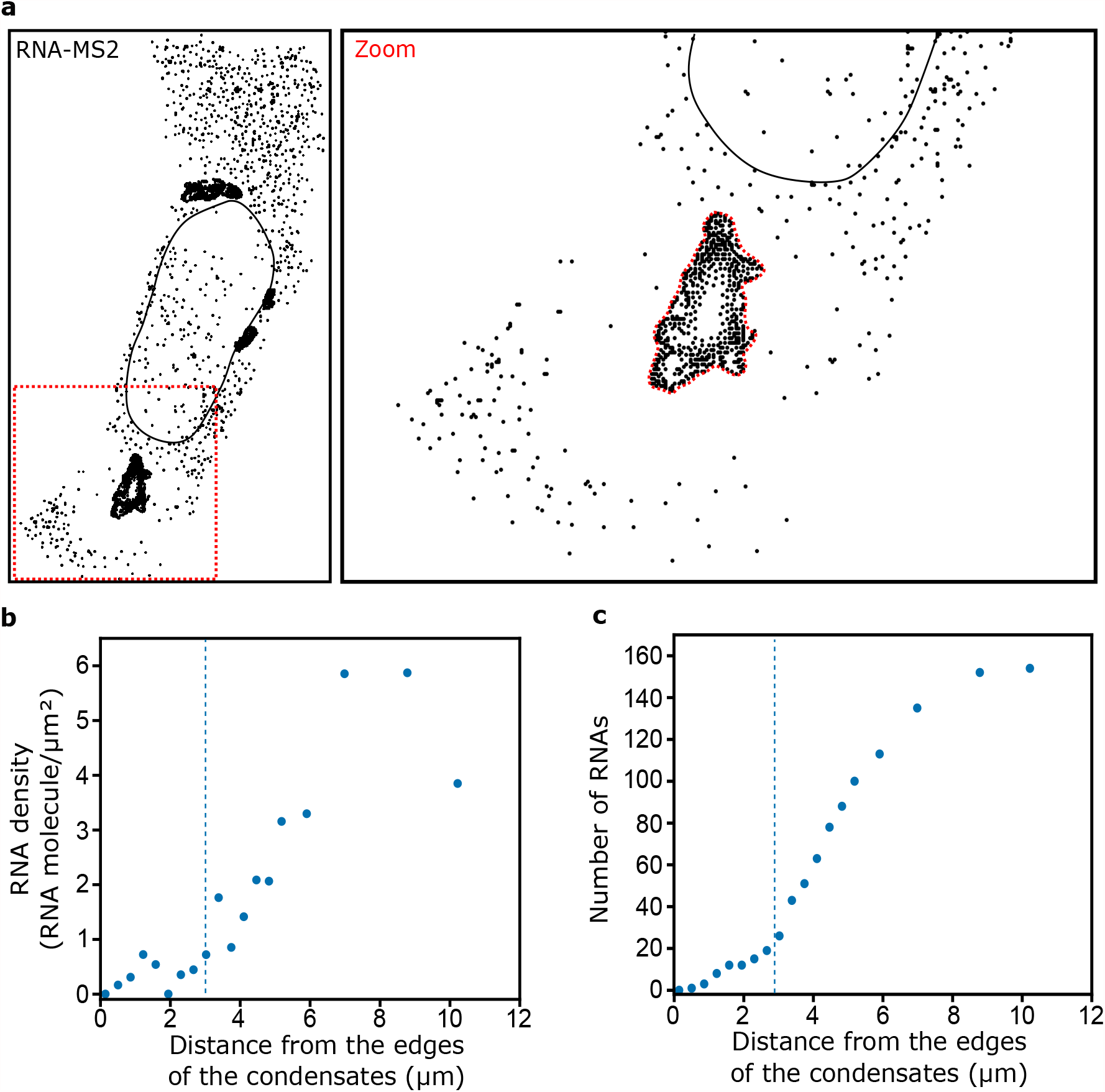
Depletion of RNA-MS2 around ArtiG^emGFP/MCP^ condensates. **a. b.** and **c.** show another example of RNA-MS2 depletion, analysed as in figure **2a. c. d.**

**Figure S3:**
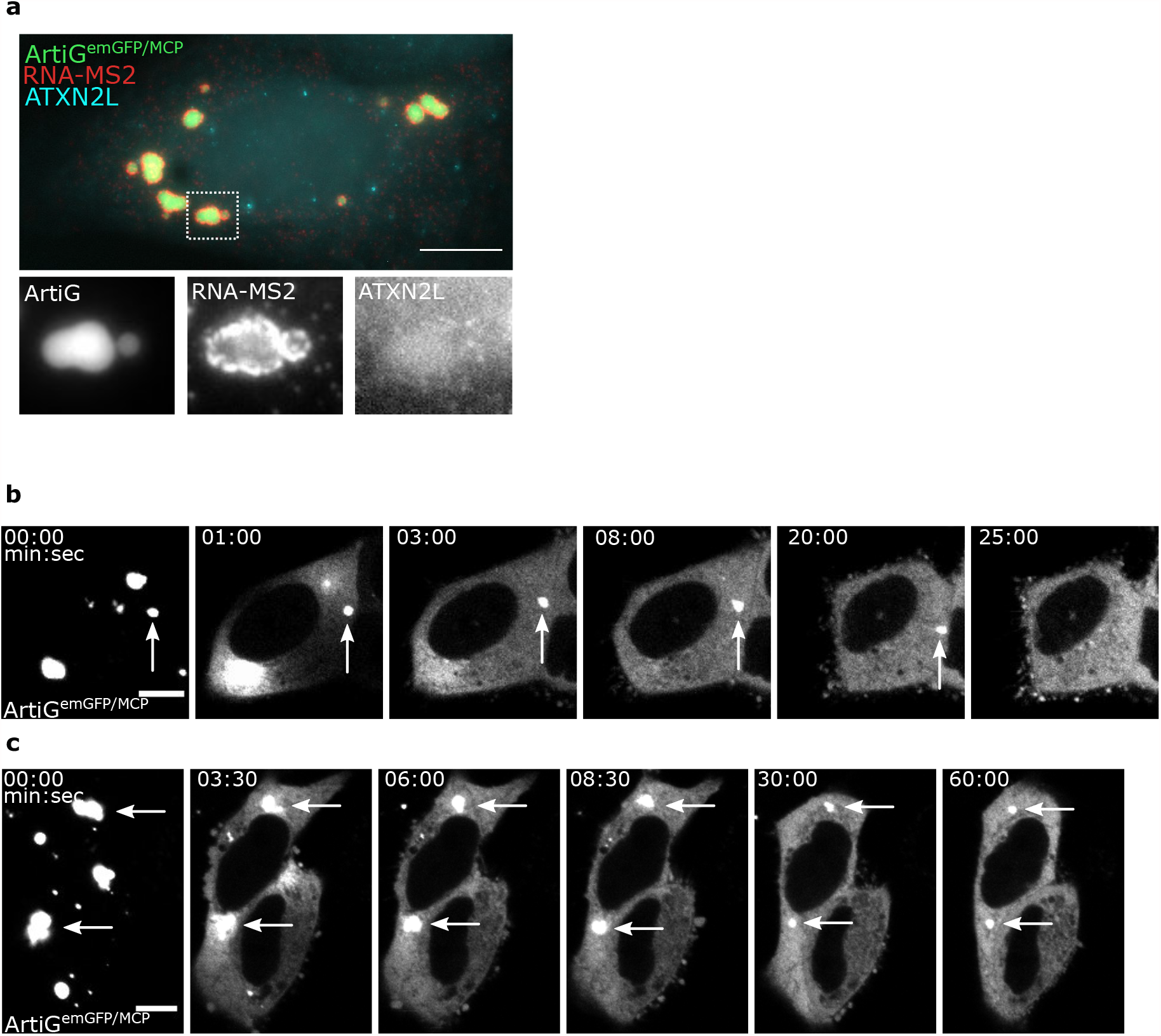
ArtiG^emGFP/MCP^ do not induce SGs and dissolve upon FK506 treatment. **a.** Epifluorescence imaging of ArtiG^emGFP/MCP^ (green) and RNA-MS2 (red) in HeLa cells, after immunostaining of ATXN2L as a SG marker (no arsenite treatment). Scale bar = 10µm. **b.** Illustrative example of a slow condensate dissolution, which completed after 25min FK506 treatment (white arrow). Scale bar = 10µm. **c.** Illustrative example of partial condensate dissolution, which was still incomplete after 60min FK506 treatment (white arrow). Scale bar = 10µm.

**Figure S4:**
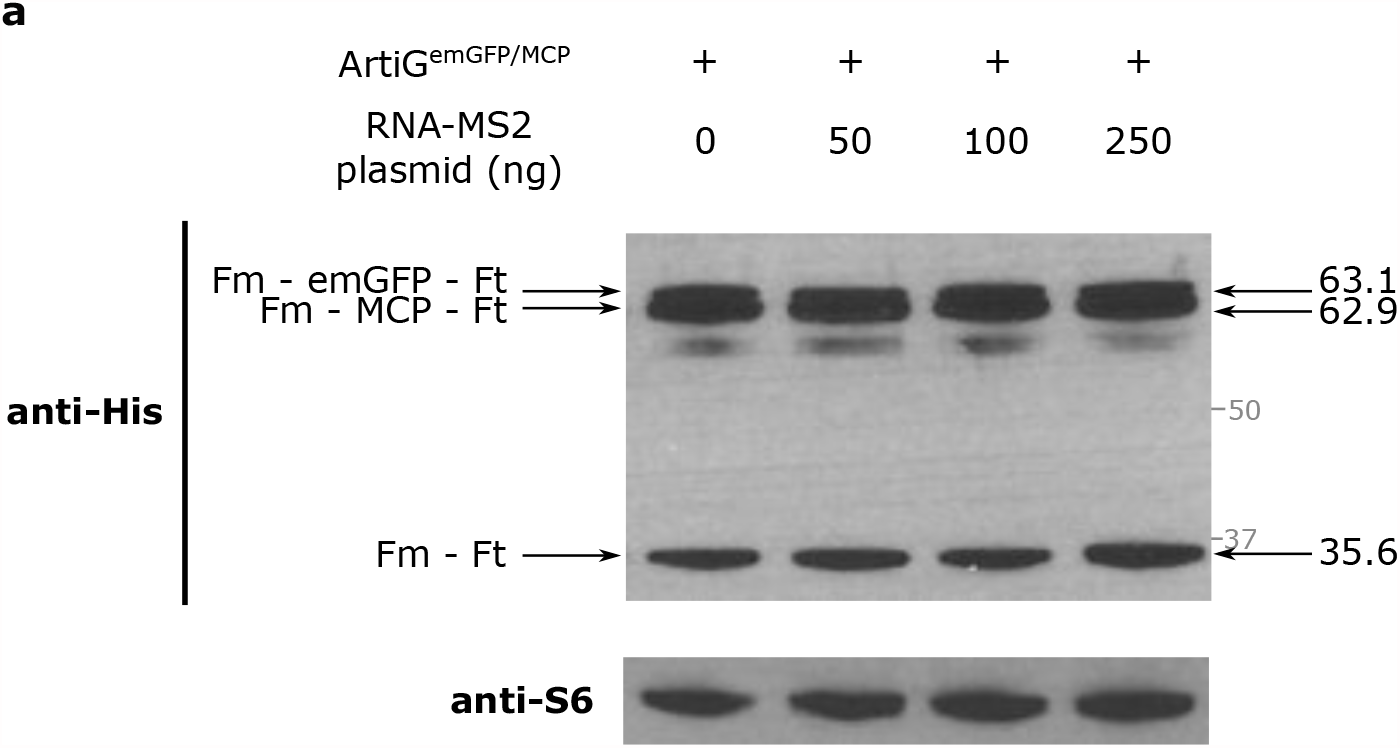
Increasing the quantity of transfected RNA-MS2 plasmid has no impact on the expression of ArtiG^emGFP/MCP^ scaffold proteins. **a.** Western blot showing scaffold protein expression 24h after transfection with Fm–emGFP–Ft, Fm–MCP–Ft and Fm–Ft and increasing amount of RNA-MS2 plasmid. The position of MW markers (in kD) is indicated on the right in grey.

**Supplementary Table 1.**
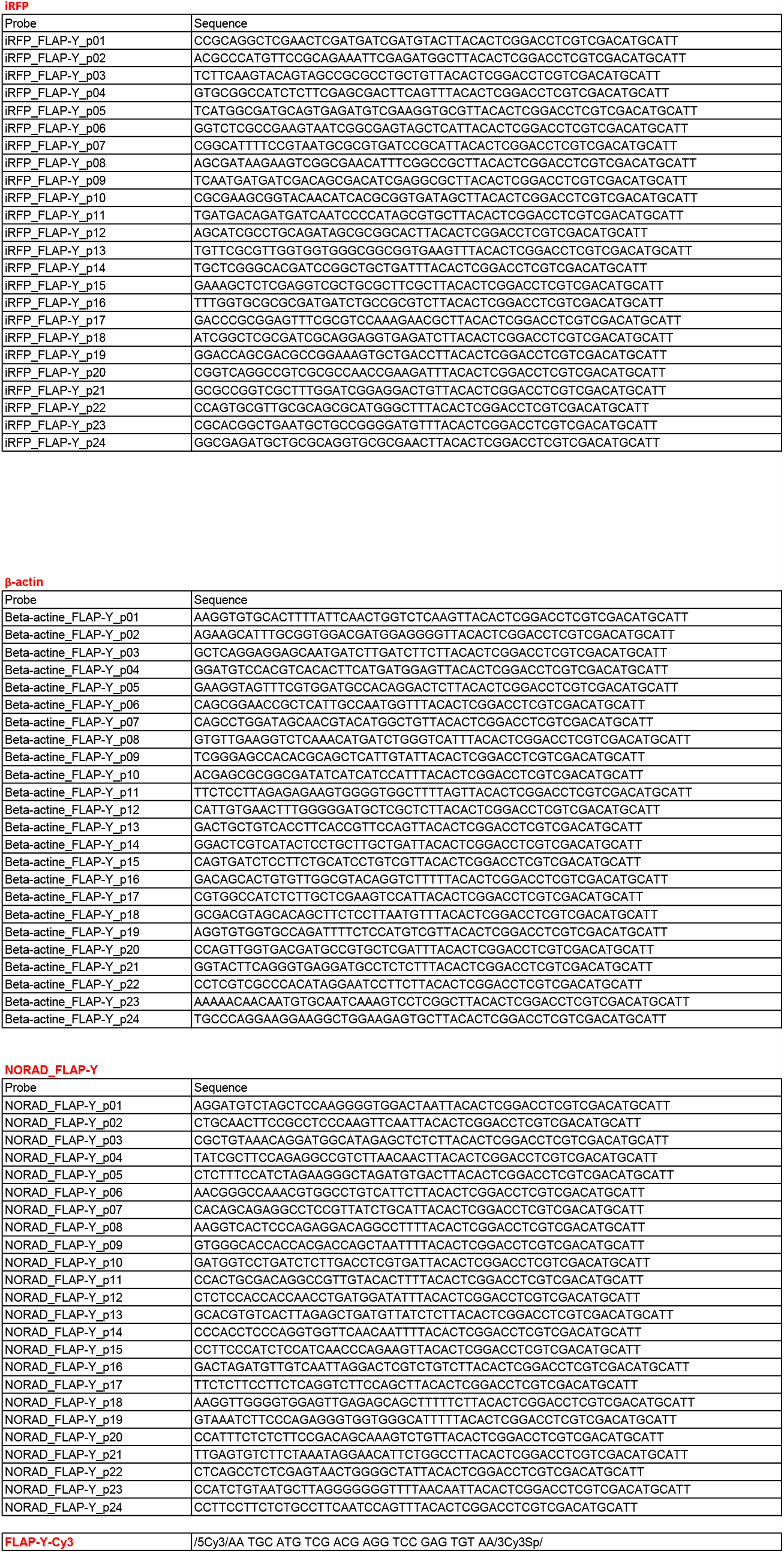

## Notes

### Competing Interest Statement

The authors have declared no competing interest.

